# Female Zebrafish (*Danio rerio*) Demonstrate Stronger Preference for Established Shoals over Newly-Formed Shoals in the Three-Tank Open-Swim Preference Test

**DOI:** 10.1101/2022.03.07.483359

**Authors:** Andrew J. Velkey, Caroline H. Koon, Isabel A. Danstrom, Katie M. Wiens

## Abstract

Zebrafish (*Danio rerio*) share a considerable amount of biological similarity with mammals, including identical or homologous gene expression pathways, neurotransmitters, hormones, and cellular receptors. Zebrafish also display complex social behaviors like shoaling and schooling, making them an attractive model for investigating normal social behavior as well as exploring impaired social function conditions such as autism spectrum disorders. Newly-formed and established shoals exhibit distinct behavior patterns and inter-member interactions that can convey the group’s social stability. We used a three-chamber open-swim preference test to determine whether individual zebrafish show a preference for an established shoal over a newly-formed shoal. Results indicated that both sexes maintained greater proximity to arena zones nearest to the established shoal stimulus. In addition, we report the novel application of Shannon entropy to discover sex differences in systematicity of responses not revealed by unit-based measurements; male subjects spent more time investigating between the two shoals than female subjects. This novel technique using established versus newly-formed shoals can be used in future studies testing transgenics and pharmacological treatments that mimic autism spectrum disorder and other disorders that affect social interaction.

## Introduction

Zebrafish share many relevant genes with mammals, promoting the species as a useful model for neuroscience, biomedical and human behavioral disorder research [1, 2]. Zebrafish have been used as an animal model for disorders of the nervous system including anxiety [1, 3], addiction [4], epilepsy [5], and several neurodegenerative diseases [6, 7]. Zebrafish also demonstrate sociability and many aspects of grouping behaviors [8-10] and are therefore a viable model for investigating social behavior as well. Shoals provide social benefits such as greater proximity with potential mates, enhanced foraging efficiency, as well as the dilution effect [11] even as social grouping has certain costs such as increased competition for limited resources and stronger parasite/disease transfer between individuals [12].

Zebrafish display a strong preference towards joining a shoal with live conspecifics versus remaining socially isolated [13, 14, 15, 16]. Group living enhances survival through multiple anti-predatory defense mechanisms. Due to the many eyes hypothesis, the predation risk of shoaling can be minimized by the benefits of aggregation, as a large group is better equipped than an isolated animal to detect predation compared [17, 18]. Heightened shoaling behavior could also be an indicator of positive affect as it can be spontaneous and involves fewer antagonistic interactions [19]. However, group living can also provide adverse circumstances, as parasites are more easily spread in large groups, and large groups can be more prone to targeting by predators [20]. In addition, intra-group competition for food and resource limitation grows as shelter size increases [12]. It is necessary to determine affinity for specific shoal characteristics to completely understand the nuances of this complex behavior and apply it to human disease models.

While many studies have addressed zebrafish preference based on visual characteristics such as fish size [21], shoal size, male-to-female ratio, stripe pattern, etc. [22], the role of intragroup familiarity in zebrafish social preference is not known. Our results indicate that a single test subject can differentiate between an established shoal and a newly-formed shoal, and both male and female subjects prefer to spend more time in proximity with the established shoal. Female test fish explore more within the vertical column nearest the established shoal while male test fish make more cross-tank transitions.

### Zebrafish exhibit preferences in social choice

Previous studies indicate characteristics driving fish shoal preference such as shoal size [23], shape [24], and parasite load [25]. Visual cues specific to individual fish within shoals is also an important determinant of shoaling behavior. Zebrafish are able to identify phenotypic differences in stripe pattern and exhibit shoaling preference dependent on early exposure to specific patterns that do not depend on their own phenotype [26, 27]. European minnows prefer to interact with shoals that are known to them [28] even if the familiar shoal is the smaller shoal; however, no research has presented subjects with a choice between novel shoals that have different levels of inter-member familiarity. Social preferences based on visual cues suggest that shoal features, like inter-member familiarity, may also influence preference of a lone fish.

#### Sex differences for shoaling preference

Certain aspects of shoaling behavior differ between males and females [29]. Interestingly, male zebrafish display bolder responses than females in both the open-field test and the novel object test [30], and male zebrafish are more exploratory of novel environments than females [31]. Shoal sex and size represent two qualities that may influence the affinity a subject has for a particular shoal. Previous studies have supported the presence of sex differences in zebrafish involving choice between joining a shoal or remaining segregated [29]. In one study, a single subject chose to spend time in proximity with a singular same-sex fish than with a conspecific of the opposite sex [15, 31]. However, another study that observed subject partiality between shoals demonstrated that males preferred to shoal with females, though females demonstrated no clear preference to shoal with one sex over the other [15]. Though male zebrafish have demonstrated a lack of significant preference for a particular shoal size, females display a clear preference for larger shoals compared to smaller ones [15, 14]. Because females may seek shoaling for protection purposes, prioritizing shoal size over shoal sex is anticipated in female shoaling behavior; larger shoals provide greater protection from predators through enhancing the confusion effect [23, 32] and increasing the dilution effect [33, 34], thus affording greater protection for each member of the shoal [14]. Previous research indicates that predation risk is a likely reason for female sexual segregation [31]. The dynamics and safety of shoal stability may also impact a female’s shoal preference when shoal size is modulated [14]. However, both male and female zebrafish prefer to affiliate with shoals rather than remain socially isolated, emphasizing innate social tendencies commonly observed among the species [15, 14]. Because both females and males show preferences for different shoal characteristics such as size and shape, additional factors likely influence zebrafish shoaling behavior; other criteria may include parasitism [35], distance from predators [36], and presence of poor competitors [37].

#### Novel vs. established shoals

A newly formed shoal generates immediate competition between members, resulting in the formation of dominant and subordinate roles within the shoal [38]. A previous study examined the activity of both dominant and subordinate male zebrafish immediately after grouping and again after five days of acclimation to conspecifics. Behavioral observations on the fifth day suggest that the formation of a stable social hierarchy occurs within the first five days following initial group formation [39]. These visible alterations in behavioral tendencies after shoal formation suggest that an established shoal may appear as a less threatening environment, while a newly-formed shoal may present as a more harmful environment due to the lack of established hierarchical roles. In addition to known sexually dimorphic behavioral characteristics in zebrafish [40, 41], dominance and aggression behavior patterns within shoals might differ between males and females. For instance, dominant males are more aggressive with their submissive conspecifics than dominant females [42], and males demonstrate stronger lateralization during aggressive responding than females [43].

Behavioral markers of previously established shoals have been observed and characterized. The Trinidadian guppy (*Poecilia reticulata*) and fathead minnow (*Pimephales promelas*) exhibit shoaling behaviors similar to those displayed by zebrafish and are also used as models for sociality and social behavior [44, 45]. Findings from investigations on Trinidadian guppy shoal fusion indicate a gradual decrease in the mean difference in shoal member size after two shoals were introduced. Nearly all previously established shoals reformed new groups due to fish size preference [46]; the shoal fusion that took place illustrates the capacity of fish to choose a shoal based on member characteristics. Previous explorations of fathead minnows have revealed variations in behavioral responses under conditions that are indicative of a predatory threat [47]. Through observing behaviors of familiarity, or lack thereof, one can determine whether a newly-formed or established shoal is more stable when presented with a potential predator. Established shoals with intergroup familiarity demonstrated more tight shoaling behavior, less freezing, and more dashing behavior. Additionally, these members performed more inspection visits compared to subjects in the newly-formed shoal [47]. These findings support the notion that a previously established shoal provides heightened security during potential threats of predation. Based on the increased likelihood of survival, it is possible that zebrafish and minnows may choose to join a shoal that presents as established. In addition, previous research suggests that intragroup familiarity is accompanied by more efficient communication between members, providing a less dangerous social living space [48, 49].

Shoal cohesion is characterized by interactions between conspecifics in response to their changing environment [50]; therefore, observing intermember communications may help determine the capacity for different shoals to attract isolated fish. Recently, investigators examined the preference of both male and female wildtype zebrafish upon providing subjects with the choice between a familiar fish and a novel fish. Both male and female subjects exhibited a preference for a novel conspecific rather than a familiar one [13]. The implications from this study strengthen the conjecture that zebrafish possess social memory. It is therefore likely that swimming pattern and tank localization can serve as dependent variables to identify affinity toward shoals of varying levels of establishment.

We suggest that, in conjunction with the ability to differentiate between shoals, zebrafish are capable of detecting the degree of polarization of a particular shoal by observing swimming patterns of the shoal members. Previous investigations have observed longitudinal patterns of shoaling and schooling of zebrafish to identify characteristic differences of the two behaviors [51]. Polarization represents the tendency of a group to swim in the same direction. Based on differential characteristic swimming patterns, shoals are seen as a low-polarized group, while a school is considered a high-polarized group [11, 18]. When relative location and average swim speed of subjects were analyzed over a five-day period, polarization decreased after this interval, possibly due to a higher level of perceived comfort or safety [51]. These findings raise the question of whether acclimatizing to an established shoal appears more attractive to a single test fish rather than acclimatizing to a newly formed, and possibly more polarized, shoal. Namely, will a new zebrafish “fit in” better with an established shoal because it is seeking comfort and safety?

Prior research has not only considered the innate social tendency of zebrafish to shoal [29] but has also investigated numerous aspects of zebrafish behavior within a shoal [47, 11, 52]. Nonetheless, few studies have examined the role of cohesion and shoal stability in determining preferences between shoals. Further, though some studies support the conjecture that zebrafish exhibit evidence of social memory [13, 53], it is still unknown whether a lone fish can perceive the visual differences of a newly-formed versus an established shoal, and how those differences might affect social choice. Prior research in our lab demonstrated the experimental efficacy of a novel open-swim paradigm for studying zebrafish social preference [54]. We used the open-swim task to test zebrafish preference for an established shoal over a newly-formed shoal. The test fish was placed in the center compartment of a three-chamber tank system while both an established and a newly-formed shoal were each displayed in the two flanking tanks. Given the reduced aggression and increased cohesion found in established shoals, both male and female zebrafish subjects demonstrated proximal preference for an established shoal over a newly-forming shoal. Furthermore, notable sex differences were demonstrated with female zebrafish showing stronger preference for established shoals over newly-formed shoals while also displaying lower behavioral entropy than male zebrafish. The results from the current study expand the present knowledge on shoaling preference and can be used in future studies of social preference in wild-type as well as investigational lines of zebrafish.

## Method

### Subjects

The subjects (*N* = 82) were healthy mature male (*n* = 45) and female (*n* = 37) wild-type zebrafish (EKK strain) of approximately 6-12 months of age and 2.5 - 6.4 cm in length. Subjects were obtained from Aquatica Tropicals, Inc. (Ruskin, Florida, USA). All care and treatment of subjects in the present study were consistent with the recommendations in the Guide for the Care and use of Laboratory Animals [55]. The research was conducted under an existing protocol (#2019-8) reviewed and approved by the Christopher Newport University Institutional Animal Care & Use Committee.

### Materials, apparatus, and procedure

Subjects were sexed and housed in four separate holding tanks (76.2 × 31.8 × 32.4 cm), each holding 75.7 liters of conditioned water maintained at a temperature of 28.5° C. Male and female zebrafish were kept in eight separate same-sex groups for the entirety of the experiment. The water had constant filtration and aeration systems, and all fish were housed under a 14-hour light/10-hour dark cycle. Fish were fed daily using the Aquaneering Scientific Hatcheries Diet for *Danio rerio*. The present study used newly-formed and established shoals as stimuli. Novel shoals were grouped in the morning and used the same day of experimentation as a stimulus; to ensure the demonstration of behaviors that characterize a shoal as novel, shoal members were obtained from separate holding tanks. Shoals were held for seven days in groups of four fish in order to establish intra-shoal familiarity and were subsequently used as the established shoal stimulus.

The objective of the present study was to determine the behavioral responses of test subjects when they were presented with different shoaling stimuli. Experimental subjects were selected for each trial and performed an open-tank, free swim task [13, 54]. The testing tank (30.5 × 15.2x 20.3 cm) was one 7.6-liter tank positioned between two stimulus tanks of the same size (Fig 1). Rosco brand Linear Polarizing Filter Sheets (#7300) were obtained from B & H Photo and Video (New York, NY) and were placed on the outer surface of the center tank between the side tanks. One of the filters had the grid oriented horizontally and the other grid was vertically oriented. With the filters in place, the stimulus fish of the tank on one end were unable to see the stimulus fish on the opposite end, but the subject in the center tank was able to see the stimuli in each flanking tank. The two stimulus shoals each contained four zebrafish, all of the same sex as the test subject, but taken from different home tanks than the subjects. For each stimulus pairing, the position of each stimulus type was counterbalanced such that an equivalent number of trials were run with each stimulus on the left side as the number of trials with each stimulus on the right side. The two stimuli in each flanking compartment consisted of an in-sex newly-formed shoal on one side and an in-sex established shoal on the opposite side.

**Figure 1.**
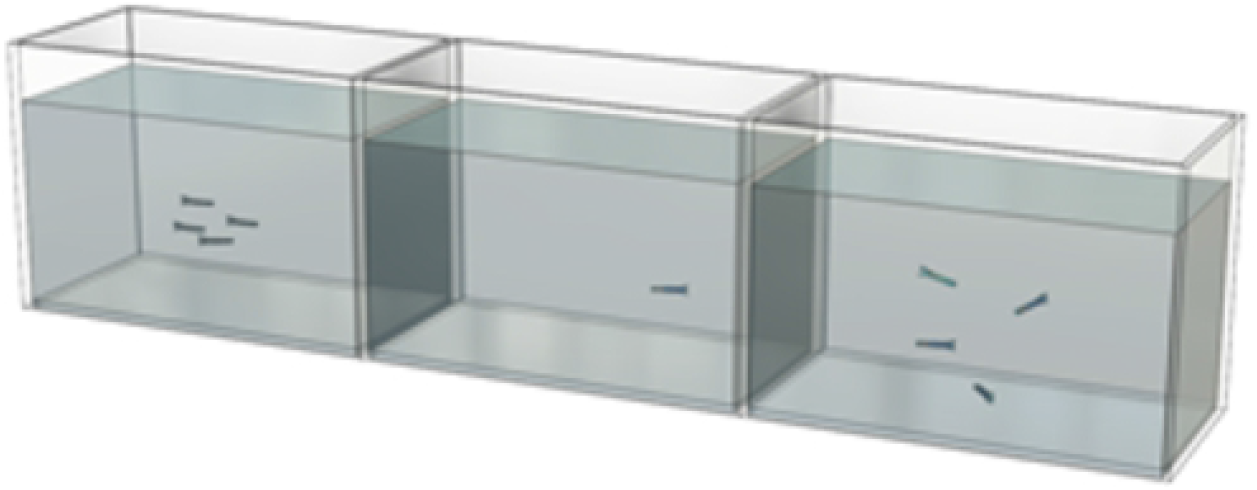
Experimental Apparatus. Illustrative example with the subject in the center tank to test preference between an established shoal on the right and a newly-formed shoal on the left. The apparatus consists of three identical 9.5 L (2.5 gal.) glass aquaria (each 30.5 cm wide × 20.3 cm high × 15.3 cm long; total width = 91.5 cm) filled with water to a depth of 16 cm.

Behavior tracking and analysis was conducted using EthoVision XT 15.0. Acquisition of tracks of the test subject via EthoVision XT 15.0 initiated after a 3-minute habituation period. Prior to and during the habituation period, the lateral sides and rearmost wall of the experimental tank were opaque in order to reduce the influence of the surrounding area. The lateral sides of the center tank were covered with removable opaque barriers to obstruct the test subjects’ vision into the flanking tanks. After the 3-minute habituation period, the partitions on either side of the central tank were removed; the rearmost wall remained opaque. During the subsequent 6-minute period of free swimming, EthoVision XT 15.0 recorded and tracked the subject’s behavior.

Subject position within the central testing tank was recorded using a digital video camera mounted on a tripod positioned directly in front of the central testing tank. EthoVision XT 15.0 is a video tracking software program that can detect an animal in a live video feed, distinguish it from the background, and track the animal’s movement, behavior, and activity. EthoVision XT 15.0 was programmed to analyze each recording in real-time, thus acquiring measurements of spatial location and subjects’ proximity to adjacent tanks. Additionally, data were extracted offline to refine evaluation of swimming patterns and density. The tracking area within the central testing tank was divided into four quadrants (upper left, lower left, upper right, and lower right of the tank). Each quadrant of the arena constituted 25% of the total arena. Zebrafish preferences were characterized by swimming patterns as the program quantified time spent in each zone as well as zone transitions [56, 54]. Spatiotemporal data for each subject were captured and processed using EthoVision XT 15.0

### Design, measures, and analyses

This experiment was a 2 (Subject Sex) X 2 (Shoal Type) mixed-factorial design which was counterbalanced across both levels of presentation side for each shoal type (Novel-Left vs Established-Right or Established-Left vs Novel-right). Because no significant main effects or interactions were found for the Side factor, data were collapsed across Side for subsequent factorial analyses. The experimental design allowed for the experimental factors to be crossed against a flexibly-defined observation zone as an additional factor. As such, the analyses included four levels of quadrant (top left, bottom left, top right, bottom right) or three levels of vertical zone (left third, middle third, right third). While most statistical analyses used quadrants as the levels for the observational zone factor, certain analyses (e.g. side preference) were better examined using vertical zones as the levels of the observational zone factor. The following measures were obtained during the session with each subject:

- Cumulative duration percent within each quadrant: EthoVision quantified the total time each subject spent in each quadrant during the entire session, and the subsequent percentage of time within each quadrant was calculated for each subject’s session.
- Percent of session time moving: EthoVision quantified the total duration of tracks recorded in each quadrant while the subject was moving at any velocity.
- Percent of session time freezing: EthoVision quantified the total duration of tracks recorded in each quadrant where the subject had ceased any detectable movement for a minimum of 3 seconds. In addition, swimming patterns were monitored by experimenters; in the case where the subject displayed no movement for one minute or longer, the trial was discontinued and data excluded from analysis.
- Average movement velocity during session: EthoVision quantified the movement speed in mm/s for each subject’s movement within each quadrant which was used to calculate the average movement velocity within each quadrant during the session.
- Variability in Velocity: The Standard Deviation in average movement velocity was calculated from the measurement of movement speed within each quadrant.
- Number of transitions between quadrants: EthoVision quantified the number of transitions between adjacent quadrants (top right to bottom right & top left to bottom left as well as top right to top left & bottom right to bottom left). Similarly, EthoVision quantified the number of transitions between vertical zones (left, center, & right).
- Cross tank transitions: EthoVision quantified the number of transitions from the right side of the arena to the left side of the arena.

Using IBM SPSS (v.26), factorial data were analyzed using a Linear Mixed Model (LMM) with Type III Sums of Squares at α = .05. As heterogeneity of variance is common with these types of data [54], the model was set with a diagonal covariance structure and degrees of freedom for the denominator were adjusted using the Maximum Likelihood estimator for the LMM. Significant main effects and interactions were explored using unplanned comparisons with Bonferroni correction for family-wise error.

In order to characterize behavioral diversity across all zones in the observational arena, a single variable index based upon Shannon entropy [57] was calculated using the following formula:

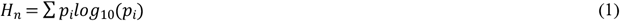

Where *H*_*n*_ is the index of behavioral diversity, *p*_*i*_ is the proportion of cumulative session time spent in zone *i*, and *n* is the total number of zones characterized with the index. The value of *H*_*n*_ can range from 0 (only systematic variability) to 1.0 (completely random variability); higher values of *H*_*n*_ indicate lower systematic variability in zone selection. This index has been used to characterize the movement of Humboldt penguins (*Spheniscus humboldti*) in a naturalistic zoo enclosure divided into zones of unequal sizes in order to examine the effects of live feeding events on the behavioral diversity of subjects across the enclosure [58]. Therefore, *H*_*n*_ is useful as a measure of characterizing systematic and random variability across a number of measures (e.g. duration in each zone) with a single index (for a review of behavioral diversity indices, see [59]). If the proportion of session time a subject in the current study spent in any particular zone is 1.0, then *H*_*n*_ = 0.0. If a subject’s proportionate time in all zones is equal across the zones, then *H*_*n*_ = 1.0. Indices of *H*_*n*_ between 0.0 and 1.0 indicate the extent to which a subject is systematically preferring any zone over the other zones.

## Results

### Duration within quadrants

Overall, subjects spent more time in lower zones adjacent to test shoals than in upper zones. Qualitatively, this difference in zone preference is demonstrated by location heat maps generated with Ethovision (see Fig 2). However, cumulative heat maps are limited in providing detail on the magnitude of these differences, which is better characterized using quantitative analyses. When analyzing the percent cumulative duration within quadrants, there was a significant main effect of Quadrant, *F* (3,128.093) = 55.336, *p ≤* .001; however, there was no significant main effect of Sex, *F* (1,230.397) = 0.562, *p* = .454, nor was there a significant interaction of Sex by Quadrant, *F* (3,128.093) = 2.081, *p* = .106 (Fig 3).

**Figure 2.**
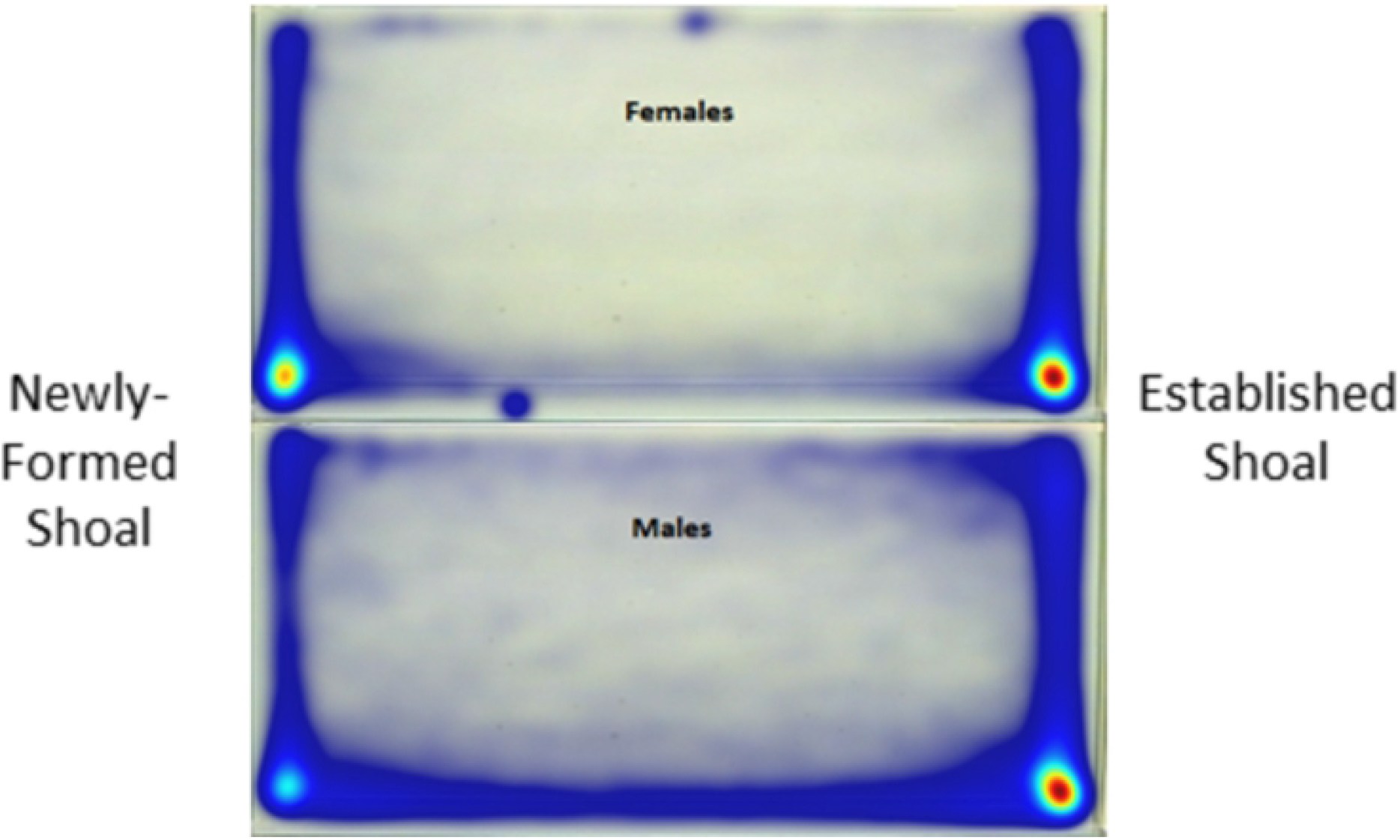
Heatmaps collapsed across trials with the same configuration of stimuli. Warm colors indicate highest intensities of localization while cooler colors indicate lowest intensities of localization.

**Figure 3.**
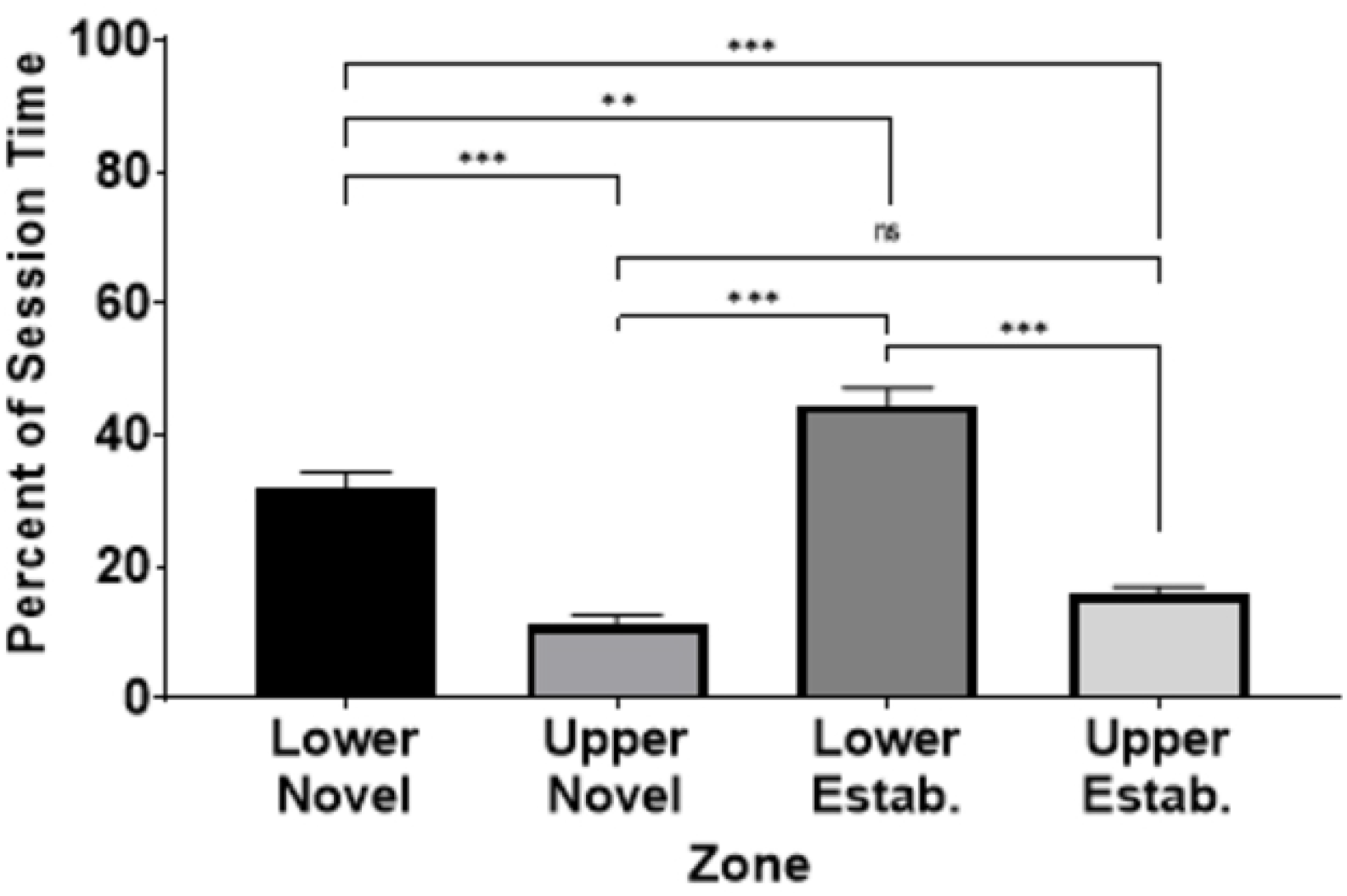
Percent cumulative duration in each quadrant zone across both sexes. Quadrants are labeled as upper/lower sections of the tank and novel/established stimulus shoal proximity. Error bars are +/- *SEM*. * indicates *p* < 0.05, ** indicates *p* < 0.01, and *** indicates *p* < 0.001.

Across both sexes, subjects spent the most time in the lower quadrant closer to the established shoal (*M* = 44.57, *SEM* = 2.618), followed by the lower quadrant closer to the novel shoal (*M* = 31.80, *SEM* = 2.524). Subjects spent less time in the upper quadrant closer to the established shoal (*M* = 15.69, *SEM* = 1.272) and the least amount of time in the upper quadrant closer to the novel shoal (*M* = 11.36, *SEM* = 1.216) (Fig. 3). The only pairwise comparison that is not significant is the difference between the time in the upper quadrant closer to the novel shoal and the upper quadrant closer to the established shoal (all other *p*’s *≤* .003).

### Time in motion

When analyzing the subjects’ percent of session time in motion within quadrants, there was a significant main effect of Quadrant, *F* (3,128.093) = 27.000, *p ≤* .001; however, there was no significant main effect of Sex, *F* (1,283.203) = 0.382, *p* = .537, nor was there a significant interaction of Sex by Quadrant, *F* (3,141.081) = 0.797, *p* = .498 (Fig 4a).

**Figure 4.**
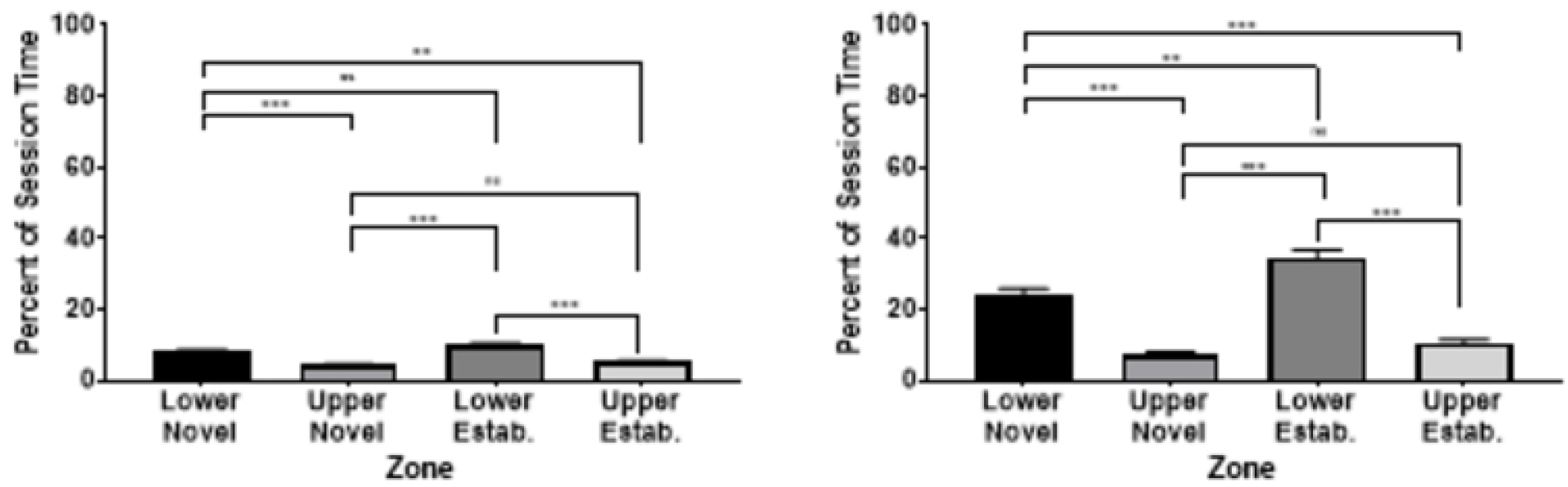
Percent duration in motion (a) and motionless (b) by zone across both sexes. Quadrants of the subject tank were labeled by upper/lower sections of the tank and novel/established stimulus shoal proximity. Error bars are +/- *SEM*. * indicates *p* < 0.05, ** indicates *p* < 0.01, and *** indicates *p* < 0.001.

Across both sexes, subjects spent the greatest percentage of time moving in the lower quadrant closer to the established shoal (*M* = 9.91, *SEM* = 0.562), followed by the lower quadrant closer to the novel shoal (*M* = 8.00, *SEM* = 0.397). Subjects spent less time moving in the upper quadrant closer to the established shoal (*M* = 5.19, *SEM* = 0.397) and the least amount of time moving in the upper quadrant closer to the novel shoal (*M* = 4.24, *SEM* = .417). The only pairwise comparisons that were not significant were the difference between the time in the upper quadrant closer to the novel shoal and the upper quadrant closer to the established shoal and the difference between the percent time moving in the lower quadrant closer to the novel shoal and the lower quadrant closer to the established shoal (all other *p*’s *≤* .001).

### Time motionless

When analyzing the percent of session time motionless within quadrants, there was a significant main effect of Quadrant, *F* (3,124.260) = 58.281, *p ≤* .001; however, there was no significant main effect of Sex, *F* (1,215.332) = 1.26, *p* = .264, nor was there a significant interaction of Sex by Quadrant, *F* (3,124.260) = 2.625, *p* = .053; (Fig 4b).

Across both sexes, subjects spent the most time motionless in the lower quadrant closer to the established shoal (*M* = 34.66, *SEM* = 2.192), followed by the lower quadrant closer to the novel shoal (*M* = 23.79, *SEM* = 2.524). Subjects spent less time in the upper quadrant closer to the established shoal (*M* = 10.50, *SEM* = 0.943) and the least amount of time in the upper quadrant closer to the novel shoal (*M* = 7.12, *SEM* = 0.865). The only pairwise comparison that is not significant is the difference between the time in the upper quadrant closer to the novel shoal and the upper quadrant closer to the established shoal (all other *p*’s *≤* .002).

### Average velocity

When analyzing the subjects’ average velocity within quadrants, there were significant main effects of Quadrant, *F* (3,180.743) = 22.925, *p ≤* .001, and Sex, *F* (1,282.043) = 6.398, *p* = .012 (Fig 5a); however, there was no significant interaction of Sex by Quadrant, *F* (3,180.743) = 0.582, *p* = .628; Females (*M* = 60.26 mm/s, *SEM* = 2.503) had a faster average velocity than males (*M* = 51.84 mm/s, *SEM* = 2.195)..

**Figure 5.**
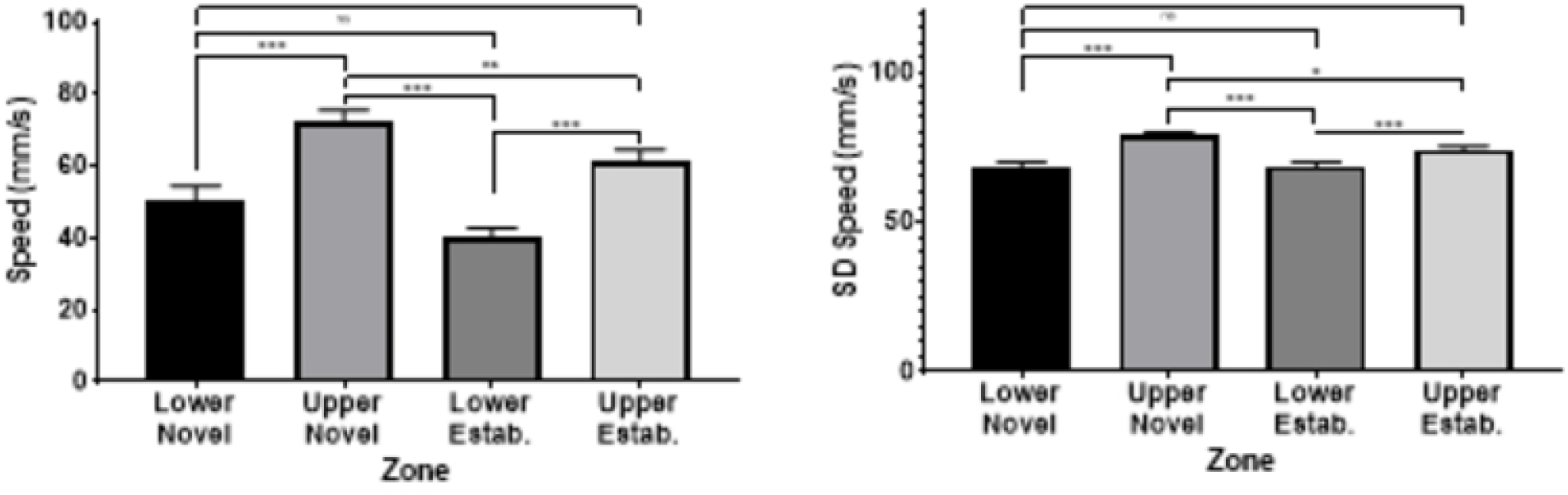
Average velocity (a) and velocity SD (b) during movement. Quadrants of the subject tank were labeled by upper/lower sections of the tank and novel/established stimulus shoal proximity. Error bars are +/- *SEM*. * indicates *p* < 0.05, ** indicates *p* < 0.01, and *** indicates *p* < 0.001.

Across both sexes, subjects moved at the slowest velocity in the lower quadrant closer to the established shoal (*M* = 40.34 mm/s, *SEM* = 2.342), followed by the lower quadrant closer to the novel shoal (*M* =50.52, *SEM* = 4.028). Subjects swam faster in the upper quadrant closer to the established shoal (*M* = 61.20 mm/s, *SEM* = 3.413) and swam the fastest in the upper quadrant closer to the novel shoal (*M* = 72.15 mm/s, *SEM* = 3.313). The only pairwise comparisons that were not significant were the difference between the average velocity in the upper quadrant closer to the novel shoal and the upper quadrant closer to the established shoal and the difference between the average velocity in the upper quadrant closer to the established shoal and the lower quadrant closer to the novel shoal (all other *p*’s *≤* .001).

### Variability in velocity

One measure of variability in velocity is the standard deviation of each subject’s swim speed within each quadrant. When analyzing the subjects’ standard deviation of velocity within quadrants, there was a significant main effect of Quadrant, *F* (3,160.255) = 25.926, *p ≤* .001; however, there was no significant main effect of Sex, *F* (1,296.650) = 0.38, *p* = .847, nor was there a significant interaction of Sex by Quadrant, *F* (3,160.255) = 1.291, *p* = .279 (Fig 5b).

Across both sexes, subjects had the lowest variability in velocity while swimming in the lower quadrant closer to the established shoal (*M* = 64.70 mm/s, *SEM* = 1.213), followed by the lower quadrant closer to the novel shoal (*M* =68.57 mm/s, *SEM* = 1.581). Subjects had greater variability in their swim speeds in the upper quadrant closer to the established shoal (*M* = 73.99 mm/s, *SEM* = 1.325) and the highest variability in the upper quadrant closer to the novel shoal (*M* = 78.71 mm/s, *SEM* = 1.140). The only pairwise comparisons that were not significant were the differences between variability in swim speeds in upper quadrant closer to the established shoal and the lower quadrant closer to the novel shoal and the difference between the average variability of subjects’ swim speeds in the lower quadrant closer to the established shoal and the lower quadrant closer to the novel shoal (all other *p*’s *≤* .046).

### Zone and Cross-Tank Transitions

Inter-quadrant transitions were analyzed by a 4 (Quadrant of Origin) by 2 (Sex) LMM. There was a significant main effect for Sex, *F* (1, 312.96) = 4.63, *p* = 0.032. There was neither a significant main effect for Quadrant of Origin, *F* (1, 152.04) = 0.996, *p* = 0.397, nor a significant interaction of Quadrant of Origin by Sex, *F* (1, 152.04) = 0.236, *p* = 0.871 (Fig 6). Across all Quadrants of Origin, females (*M* = 15.55, *SEM* = 1.064) had significantly more quadrant-to-quadrant transitions than males (*M* = 12.51, *SEM* = 0.933). To assess cross-tank transitions, data were collapsed across upper and lower quadrants, and an independent-samples *t-*test was performed between males and females on cross-tank transitions. Males (*M* = 14.33, *SEM* = 1.253) had significantly more cross-tank transitions than females (*M* = 7.73, *SEM* = 1.053), *t* (80) = 3.930, *p ≤* .001.

**Figure 6.**
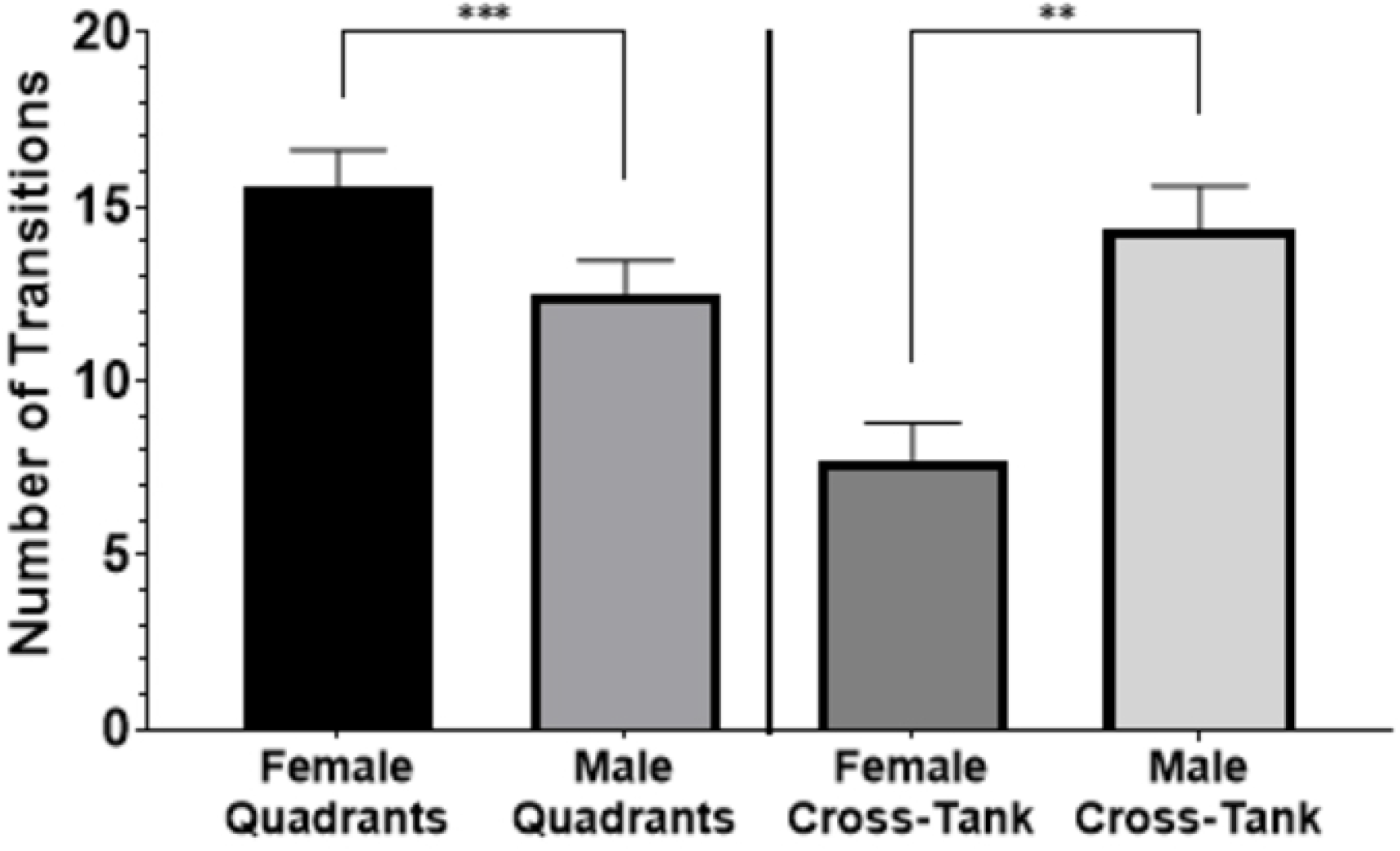
Sex differences in inter-quadrant and cross-tank transitions. Quadrants of the subject tank were labeled by upper/lower sections of the tank and novel/established stimulus shoal proximity. Error bars are +/- SEM. * indicates *p* < 0.05, ** indicates *p* < 0.01, and *** indicates *p* < 0.001.

### Behavioral Diversity Index

For the current data, two indices of *H*_*n*_ were calculated for each subject. One index, *H*_*4*_, considered the four quadrants (upper right, upper left, lower right, lower left) of the observation arena while the other index, *H*_*3*_, considered the left, middle, and right vertical thirds of the observation arena. *H*_*4*_ thus provides an index which characterizes diversity in zone selection both vertically and horizontally within the observation arena, while *H*_*3*_ provides an index which characterizes diversity in zone selection horizontally across the observation arena. Higher values of either *H*_*4*_ or *H*_*3*_ indicate more random variability in a subject’s proportions in zone utilizations. Whenever a case for a subject had a proportion for any zone equal to zero, resulting in an undefined computation for the log of that value, the undefined result was set to zero and the summation for *H*_*n*_ continued across the other non-zero proportions.

As expected, males had significantly higher behavioral entropy index scores than females for *H*_*3*_ analyzed across three zones (males *M =* 0.360, *SEM =* 0.016; females *M =* 0.243, *SEM =* 0.022; *t* (80) = 4.365, *p ≤* .001, one-tailed). Males also had significantly higher behavioral entropy index scores than females for *H*_*4*_ analyzed across four zones, (males *M =* 0.465, *SEM =* 0.016; females *M =* 0.383, *SEM =* 0.021; *t* (80) = 3.12, *p* = 0.033, one-tailed) (Fig.7). Taken together, these results indicate that male subjects show less systematic variability in their utilization of zones in the observation arena than female subjects, revealing sex differences in overall responding not revealed by main effects or interactions in the Sex X Zone factorial analyses of the specific dependent measures (except for the main effect of Sex on swim velocity).

**Figure 7.**
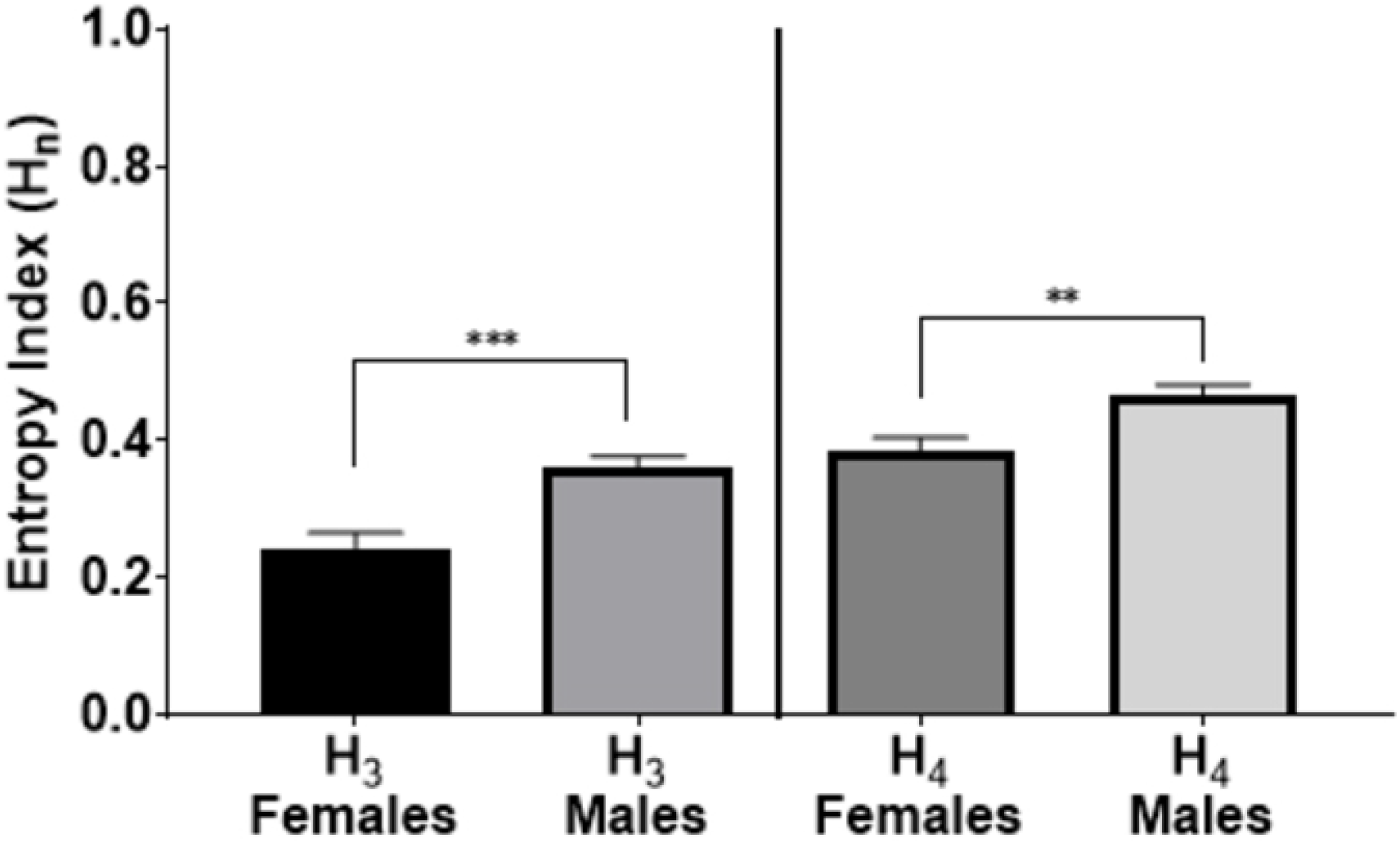
Sex differences in behavioral entropy. H_3_ measures entropy across three vertical zones while H_4_ measures entropy across four quadrants. Error bars are +/- *SEM*. * indicates *p* < 0.05, ** indicates *p* < 0.01, and *** indicates *p* < 0.001.

## DISCUSSION

The results presented here indicate that solitary zebrafish can differentiate between established and novel shoals based solely on visual cues and choose to spend more time near an established shoal. Movement measures indicate that both sexes spent more time motionless when in the lower quadrant nearest the established shoal. Velocity measures show that both sexes exhibited less variability in velocity when in the upper quadrant near the established shoal compared to the upper quadrant near the novel shoal, potentially indicative of less darting behavior. While both male and female zebrafish spent more time in the lower quadrant near the established shoal, there were significant differences in the manner by which males and females moved around the field; females appeared to investigate more locally while males investigated more across the two shoals. Use of the Shannon entropy measure further confirmed the increased behavioral entropy for males and provides a useful measure for future comparison across studies that use different methods and arenas. The addition of shoal stimuli of varying familiarity to the three-tank open-swim preference test provides an attractive and easy-to-use system for studies investigating zebrafish models of disorders that affect social behavior and recognition of social cues.

The present analyses include a novel application of Shannon entropy to characterize the diversity in zone preference in the three-tank open-swim preference test. The three-tank open-swim preference test has been used successfully in a number of previous studies, and it is becoming increasingly popular for the investigation of social preference in both normal populations (e.g. EKK or AB wild type) [54, 60, 27] and in clinical models [58, 13]. Previous studies established the efficacy of the technique and demonstrated experimental effects through the analysis of unitary measures such as duration in zone, swimming velocity, etc. The present study establishes the utility of a unitless behavioral diversity index to characterize the extent to which subjects demonstrate a systematic preference among the observational zones of interest in the three-tank open-swim preference task and allows for comparison of the diversity index across different zone characterizations (e.g. quadrants versus vertical thirds) and between subject groupings (e.g. sex). As the nature of the three-tank open-swim preference test is to determine the extent to which test subjects demonstrate a preference and/or avoidance of zones in proximity with test stimuli, the application of an index of behavioral diversity such as Shannon entropy can be useful for future studies of social preference in tasks involving movement in open arenas.

Prior studies have demonstrated a relationship between swimming speed, vertical tank location and anxiety during exposure to novel environments and other common stressors [62, 63]. Increased swimming speeds could indicate darting patterns that result from an expression of fear [64], while more time spent in the lower half of a novel tank is indicative of an anxious state which can be reversed with exposure to anxiolytic drugs [65]. Both male and female zebrafish exhibit differences in anxiety-like responses with females spending more time in the bottom half the tank during a novel tank task and more time in the dark zone during a light-dark task [66].

Further, experiments on wild-caught zebrafish show that males are bolder during feeding than females [67]. Our results show a greater percentage of time spent in the lower quadrants for both sexes, indicating an anxious state for both male and female subjects. However, the faster average swimming speeds of female subjects observed in this study could be indicative of a higher level of anxiety compared to male subjects, suggesting that the social choice paradigm can elicit subtle behavioral anxiety differences compared to the novel tank task. Further, stress levels have the potential to induce shoaling behavior as it is exhibitive of seeking out energy rationing [68]. The subtle behavioral differences exhibited by females in our study (lower entropy and higher tendency to prefer one side of the arena rather than elsewhere) is potentially due to higher stress levels.

Zebrafish place preference can be influenced by chemical [64, 69] or live stimuli [54, 70], and a three-chamber apparatus is commonly utilized to quantify preference depending on subject swimming patterns [54, 13]. As an example of place bias indicating social choice, Social Preference Index has been utilized in several studies concerning social preference in zebrafish, equating proximity with preference for interaction [71, 72, 73]. Swimming patterns localized near one stimulus over another could indicate preference, but other driving forces of social behavior, such as investigative or aggressive interactions, should not be discounted. The present study solely investigated social interaction based on visual information and did not delineate between aggression versus investigation. However, aggressive interaction or investigation would also indicate a change in social interaction that occurs due to the observed differences in stimuli shoal cohesion, suggesting that the subject fish were still able to differentiate between shoal types.

A salient component of Fetal Alcohol Spectrum Disorders (FASD), Autism Spectrum Disorders (ASD), and several other neurodevelopmental disorders’ symptomatology is the display of atypical social behavior. Since the zebrafish model exists at the intersection of behavioral complexity and biological simplicity, use of the zebrafish to study neurodevelopmental disorders has recently gained popularity. Shoaling tendencies are first distinguishably exhibited by young zebrafish approximately two weeks post-hatch, and shoaling as a species-specific behavior is critically influenced by early life experience [74]. Zebrafish exposed to alcohol at the embryonic stage have been shown to display impaired shoaling behavior development [75]. In humans, FASD is a life-long disorder, and social impairments including social withdrawal and depression persist for the duration of a patient’s life. Similarly, embryonic zebrafish exposed to ethanol exhibit severely reduced shoaling responses that continue two years following initial exposure and derive from central nervous system changes rather than motor or visual dysfunction [76]. The established versus new shoaling model outlined here contributes to understanding social behavior in typically-developing zebrafish, though the model may be useful to better classify social impairment in neurodevelopmental disorders such as FASD or ASD. Social interaction difficulties in FASD and ASD may be evident through a failure to recognize differences in established versus new shoals. Such behavior would be supported by a lack of preference for either shoal, shown by a subject spending equal amounts of time in close proximity to either stimuli.

The DYRK1A gene, located in the “Down Syndrome Critical Region DSCR” has been identified as a significant element in the pathogenesis of ASD in humans [77]. DYRK1A mutation in humans is connected to intellectual impediments, microcephaly, and ASD. When DYRK1A is knocked-out in zebrafish (DYRK1A KO), affected subjects exhibit social abnormalities parallel to those displayed by human ASD patients. Specifically, DYRK1A KO resulted in decreased expression of c-fos, a proto-oncogene important for cellular proliferation and differentiation [77, 78]. When presented with a three-member social stimuli shoal, DYRK1A KO zebrafish spent significantly less time in the zone of closest proximity to the shoal compared to wildtype (WT) zebrafish [77]. Using the DYRK1A KO in our three-chamber social choice model with established versus newly-formed shoals as flanking stimuli would determine if DYRK1A expression is necessary for identifying intragroup familiarity between shoals.

Our findings and conclusions present implications for both basic research on the mechanisms of social preference in animals as well as the aforementioned zebrafish models of human disease and behavioral dysfunction. Future research on social preference using the three-tank open-swim preference test could explore the various characteristics of intra-shoal activity that indicate social novelty within a newly-established shoal which are subsequently detected by the observing subject. In addition, future research could also explore whether other factors such as age, size, or health status affect preference for an established shoal, and whether other features of test shoals and/or individual subjects can override the preference for established shoals. In order to extend and further explore various aspects of subjects’ movement, future researchers could use a more-sophisticated dual-camera setup that allows for analysis of three-dimensional movement data [79, 80] which may subsequently reveal other, perhaps more subtle, differences in shoal preference. Finally, the demonstrated value of Shannon entropy in the current study introduces new possibilities for the comparison of arena-based movement of subjects under a variety of conditions across studies.

## Acknowledgements

The authors thank John Velkey of the University of Maryland – Baltimore County for his assistance with figures and Jake Boles of the University of Florida for technical assistance with the use of EthoVision XT 15.0. We are grateful for the Christopher Newport University Office for Undergraduate Research and Creative Activity, which provided financial support of this work through the CNU Summer Scholars Program and the Undergraduate Research Captains Program. These programs provided housing and stipends to the second author (CHK) in support of pilot work, data collection, and manuscript preparation.

## Supporting Information

**S1 File. Novel-ExpShoalSSP_PlosONE.xlsx**

## References

1. Stewart AM, Braubach O, Spitsbergen J, Gerlai R, Kalueff AV. Zebrafish models for translational neuroscience research: from tank to bedside. Trends Neurosci. 2014 May 1;37(5):264–78. doi.org/10.1152/ajpregu.00671.2007

2. Alsop D, Vijayan MM. Development of the corticosteroid stress axis and receptor expression in zebrafish. Am J Physiol Regul Integr Comp Physiol. 2008 Mar;294(3):R711–9.. doi.org/10.1016/j.tins.2014.02.011

3. Egan RJ, Bergner CL, Hart PC, Cachat JM, Canavello PR, Elegante MF, Elkhayat SI, Bartels BK, Tien AK, Tien DH, Mohnot S. Understanding behavioral and physiological phenotypes of stress and anxiety in zebrafish. Behav Brain Res. 2009 Dec 14;205(1):38–44. doi.org/10.1016/j.bbr.2009.06.022

4. Mathur P, Guo S. Use of zebrafish as a model to understand mechanisms of addiction and complex neurobehavioral phenotypes. Neurobiol Dis. 2010 Oct 1;40(1):66–72. doi.org/10.1016/j.nbd.2010.05.016

5. Baxendale S, Holdsworth CJ, Meza Santoscoy PL, Harrison MR, Fox J, Parkin CA, Ingham PW, Cunliffe VT. Identification of compounds with anti-convulsant properties in a zebrafish model of epileptic seizures. Dis Model Mech. 2012 Nov;5(6):773–84. doi.org/doi:10.1242/dmm.010090

6. Best JD, Alderton WK. Zebrafish: An in vivo model for the study of neurological diseases. Neuropsychiatr Dis Treat. 2008 Jun;4(3):567. doi.org/10.2147/NDT.S2056

7. Panula P, Sallinen V, Sundvik M, Kolehmainen J, Torkko V, Tiittula A, Moshnyakov M, Podlasz P. Modulatory neurotransmitter systems and behavior: towards zebrafish models of neurodegenerative diseases. Zebrafish. 2006 Jun 1;3(2):235–47. doi.org/10.1089/zeb.2006.3.235

8. Maaswinkel H, Zhu L, Weng W. Assessing social engagement in heterogeneous groups of zebrafish: a new paradigm for autism-like behavioral responses. PLoS One. 2013 Oct 8;8(10):e75955. doi.org/10.1371/journal.pone.0075955

9. Séguret A, Collignon B, Cazenille L, Chemtob Y, Halloy J. Loose social organisation of ab strain zebrafish groups in a two-patch environment. PLoS One. 2019 Feb 8;14(2):e0206193. doi.org/10.1371/journal.pone.0206193

10. Sykes DJ, Suriyampola PS, Martins EP. Recent experience impacts social behavior in a novel context by adult zebrafish (Danio rerio). PLoS One. 2018 Oct 18;13(10):e0204994. doi.org/10.1371/journal.pone.0204994

11. Miller NY, Gerlai R. Shoaling in Zebrafish: What We Don’t Know. Rev Neurosci. 2011;22(1):17–25.. doi.org/10.1515/rns.2011.004

12. Majeed NA, Cyriac VP, Kodandaramaiah U. To group or not to group: Effect of prolonged exposure to predation and resource availability on the shoaling behaviour of zebrafish (Danio rerio). bioRxiv. 2020 Jan 1. doi.org/10.1101/2020.07.02.183897

13. Ariyasiri K, Choi TI, Kim OH, Hong TI, Gerlai R, Kim CH. Pharmacological (ethanol) and mutation (sam2 KO) induced impairment of novelty preference in zebrafish quantified using a new three-chamber social choice task. Prog Neuropsychopharmacol Biol Psychiatry. 2019 Jan 10;88:53–65. doi.org/10.1016/j.pnpbp.2018.06.009

14. Etinger A, Lebron J, Palestis BG. Sex-assortative shoaling in zebrafish (Danio rerio). Bios. 2009 Jul;80(4):153–8. doi.org/10.1893/011.080.0402

15. Ruhl N, McRobert SP. The effect of sex and shoal size on shoaling behaviour in Danio rerio. J Fish Biol. 2005 Nov;67(5):1318–26. doi.org/10.1111/j.0022-1112.2005.00826.x

16. Snekser JL, McRobert SP, Murphy CE, Clotfelter ED. Aggregation behavior in wildtype and transgenic zebrafish. Ethology. 2006 Feb;112(2):181–7. doi.org/10.1111/j.1439-0310.2006.01139.x

17. Pitcher TJ. The behavior of teleost fishes. Boston, MA: Springer; 1986. Chapter 12, Functions of shoaling behaviour in teleosts; p. 294–337. doi.org/10.1007/978-1-4684-8261-4_12

18. Mayer PC. Economic models of fish shoal (school) size: a near comprehensive view of single species shoaling strategy. J Bioeconomics. 2010 Jul;12(2):119–43. doi.org/10.1007/s10818-010-9084-7.

19. Franks B, Graham C, Von Keyserlingk MA. Is heightened-shoaling a good candidate for positive emotional behavior in zebrafish?. Animals. 2018 Sep;8(9):152. doi:10.3390/ani8090152

20. Krause J, Ruxton GD, Ruxton G, Ruxton IG. Living in groups. Oxford University Press; 2002 Oct 10.

21. Hoare DJ, Krause J, Peuhkuri N, Godin JG. Body size and shoaling in fish. J Fish Biol. 2000 Dec;57(6):1351–66. doi.org/10.1111/j.1095-8649.2000.tb02217.x

22. Gerlai R. Animated images in the analysis of zebrafish behavior. Curr Zool. 2017 Feb 1;63(1):35–44. doi:10.1093/cz/zow077

23. Pritchard VL, Lawrence J, Butlin RK, Krause J. Shoal choice in zebrafish, Danio rerio: the influence of shoal size and activity. Anim Behav. 2001 Dec 1;62(6):1085–8.doi.org/10.1006/anbe.2001.1858

24. Ward AJ, Kent MI, Webster MM. Social recognition and social attraction in group-living fishes. Frontiers in Ecology and Evolution. 2020 Feb 4;8:15. doi.org/10.3389/fevo.2020.00015

25. Krause J, Godin JG. Influence of parasitism on shoal choice in the banded killifish (Fundulus diaphanus, Teleostei, Cyprinodontidae). Ethology. 1996 Jan 12;102(1):40–9. doi.org/10.1111/j.1439-0310.1996.tb01102.x

26. Spence R, Smith C. The role of early learning in determining shoaling preferences based on visual cues in the zebrafish, Danio rerio. Ethology. 2007 Jan;113(1):62–7. doi.org/10.1111/j.1439-0310.2006.01295.x

27. Engeszer RE, Da Barbiano LA, Ryan MJ, Parichy DM. Timing and plasticity of shoaling behaviour in the zebrafish, Danio rerio. Anim Behav. 2007 Nov 1;74(5):1269–75. doi:10.1016/j.anbehav.2007.01.032

28. Barber I, Wright HA. How strong are familiarity preferences in shoaling fish? Anim Behav. 2001 May 1;61(5):975–9. doi.org/10.1006/anbe.2000.1665

29. Snekser JL, Ruhl N, Bauer K, McRobert SP. The Influence of Sex and Phenotype on Shoaling Decisions in Zebrafish. Int J Comp Psychol. 2010 Mar 1;23(1). escholarship.org/uc/item/8n68z3pf.

30. Dahlbom SJ, Lagman D, Lundstedt-Enkel K, Sundström LF, Winberg S. Boldness predicts social status in zebrafish (Danio rerio). PLoS One. 2011 Aug 17;6(8):e23565. doi.org/10.1371/journal.pone.0023565

31. Reolon GK, de Melo GM, da Rosa JG, Barcellos LJ, Bonan CD. Sex and the housing: Effects on behavior, cortisol levels and weight in zebrafish. Behav Brain Res. 2018 Jan 15;336:85–92. doi.org/10.1016/j.bbr.2017.08.006.

31. Croft DP, Morrell LJ, Wade AS, Piyapong C, Ioannou CC, Dyer JR, Chapman BB, Wong Y, Krause J. Predation risk as a driving force for sexual segregation: a cross-population comparison. Am Nat. 2006 Jun;167(6):867–78. doi.org/10.1086/504853

32. Krause J, Ruxton GD, Rubenstein D. Is there always an influence of shoal size on predator hunting success?. J Fish Biol. 1998 Mar;52(3):494–501. doi.org/10.1111/j.1095-8649.1998.tb02012.x

33. Lehtonen J, Jaatinen K. Safety in numbers: the dilution effect and other drivers of group life in the face of danger. Behav Ecol Sociobiol. 2016 Apr 1;70(4):449–58. doi.org/10.1007/s00265-016-2075-5

34. Frommen JG, Hiermes M, Bakker TC. Disentangling the effects of group size and density on shoaling decisions of three-spined sticklebacks (Gasterosteus aculeatus). Behav Ecol Sociobiol. 2009 Jun;63(8):1141–8. doi.org/10.1007/s00265-009-0767-9

35. Barber I, Downey LC, Braithwaite VA. Parasitism, oddity and the mechanism of shoal choice. J Fish Biol. 1998 Dec;53(6):1365–8. doi.org/10.1111/j.1095-8649.1998.tb00256

36. Ashley EJ, Kats LB, Wolfe JW. Balancing trade-offs between risk and changing shoal size in northern red-belly dace (Phoxinus eos). Copeia. 1993 May 3;1993(2):540–2.doi.org/10.2307/1447157

37. Metcalfe NB, Thomson BC. Fish recognize and prefer to shoal with poor competitors. Proc R Soc Lond B Biol Sci. 1995 Feb 22;259(1355):207–10. doi.org/10.1098/rspb.1995.0030

38. Teles MC, Oliveira RF. Zebrafish: Methods and protocols. New York: Human Press; 2016. Quantifying aggressive behavior in zebrafish; p. 293–305. doi.org/10.1007/978-1-4939-3771-4_20

39. Larson ET, O’Malley DM, Melloni Jr RH. Aggression and vasotocin are associated with dominant–subordinate relationships in zebrafish. Behav Brain Res. 2006 Feb 15;167(1):94–102. doi.org/10.1016/j.bbr.2005.08.020

40. Santos EM, Kille P, Workman VL, Paull GC, Tyler CR. Sexually dimorphic gene expression in the brains of mature zebrafish. Comp Biochem Physiol A Mol Integr Physiol. 2008 Mar 1;149(3):314–24. doi.org/10.1016/j.cbpa.2008.01.010

41. Sreenivasan R, Cai M, Bartfai R, Wang X, Christoffels A, Orban L. Transcriptomic analyses reveal novel genes with sexually dimorphic expression in the zebrafish gonad and brain. PLoS One. 2008 Mar 12;3(3):e1791. doi.org/10.1371/journal.pone.0001791

42. Paull GC, Filby AL, Giddins HG, Coe TS, Hamilton PB, Tyler CR. Dominance hierarchies in zebrafish (Danio rerio) and their relationship with reproductive success. Zebrafish. 2010 Mar 1;7(1):109–17. doi.org/10.1089/zeb.2009.0618

43. Ariyomo TO, Watt PJ. Aggression and sex differences in lateralization in the zebrafish. Anim Behav. 2013 Sep 1;86(3):617–22. doi.org/10.1016/j.anbehav.2013.06.019

44. Dugatkin LA, Godin JG. Predator inspection, shoaling and foraging under predation hazard in the Trinidadian guppy, Poecilia reticulata. Environ Biol Fishes. 1992 Jul;34(3):265–76. doi.org/10.1007/BF00004773

45. Hager MC, Helfman GS. Safety in numbers: shoal size choice by minnows under predatory threat. Behav Ecol Sociobiol. 1991 Nov;29(4):271–6. doi.org/10.1007/BF00163984

46. Croft DP, Arrowsmith BJ, Bielby J, Skinner K, White E, Couzin ID, Magurran AE, Ramnarine I, Krause J. Mechanisms underlying shoal composition in the Trinidadian guppy, Poecilia reticulata. Oikos. 2003 Mar;100(3):429–38. doi.org/10.1034/j.1600-0706.2003.12023.x

47. Chivers DP, Brown GE, Smith RJ. Familiarity and shoal cohesion in fathead minnows (Pimephales promelas): implications for antipredator behaviour. Can J Zool. 1995 May 1;73(5):955–60. doi.org/10.1139/z95-111

48. Atton N, Galef BJ, Hoppitt W, Webster MM, Laland KN. Familiarity affects social network structure and discovery of prey patch locations in foraging stickleback shoals. Proc R Soc Lond B Biol Sci. 2014 Aug 22;281(1789):20140579. doi.org/10.1098/rspb.2014.0579

49. Lemasson B, Tanner C, Woodley C, Threadgill T, Qarqish S, Smith D. Motion cues tune social influence in shoaling fish. Sci Rep. 2018 Jun 28;8(1):1-0. | doi:10.1038/s41598-018-27807-1

50. Butail S, Bartolini T, Porfiri M. Collective response of zebrafish shoals to a free-swimming robotic fish. PLoS One. 2013 Oct 16;8(10):e76123. doi.org/10.1371/journal.pone.0076123

51. Miller N, Gerlai R. From schooling to shoaling: patterns of collective motion in zebrafish (Danio rerio). PLoS One. 2012 Nov 14;7(11):e48865. doi.org/10.1371/journal.pone.0048865

52. Herbert-Read JE, Perna A, Mann RP, Schaerf TM, Sumpter DJ, Ward AJ. Inferring the rules of interaction of shoaling fish. Proc Natl Acad Sci. 2011 Nov 15;108(46):18726–31. doi.org/10.1073/pnas.1109355108

53. Madeira N, Oliveira RF. Long-term social recognition memory in zebrafish. Zebrafish. 2017 Aug 1;14(4):305–10. doi.org/10.1089/zeb.2017.1430

54. Velkey AJ, Boles J, Betts TK, Kay H, Henenlotter R, Wiens KM. High fidelity: Assessing zebrafish (Danio rerio) responses to social stimuli across several levels of realism. Behav Proc. 2019 Jul 1;164:100–8. doi.org/10.1016/j.beproc.2019.04.012

55. National Research Council. Guide for the care and use of laboratory animals. 8th Ed. Washington, DC: The National Academies Press; 2011. doi.org/10.17226/12910

56. Stewart A, Gaikwad S, Kyzar E, Green J, Roth A, Kalueff AV. Modeling anxiety using adult zebrafish: a conceptual review. Neuropharmacology. 2012 Jan 1;62(1):135–43. doi.org/10.1016/j.neuropharm.2011.07.037

57. Shannon CE. A mathematical theory of communication. Bell Syst Tech J. 1948 Jul;27(3):379–423. doi.org/10.1002/j.1538-7305.1948.tb01338.x

58. Fernandez EJ, Myers M, Hawkes NC. The Effects of Live Feeding on Swimming Activity and Exhibit Use in Zoo Humboldt Penguins (Spheniscus humboldti). J Zoo Bot Gar. 2021 Mar;2(1):88–100. doi.org/10.3390/jzbg2010007

59. Miller LJ, Vicino GA, Sheftel J, Lauderdale LK. Behavioral diversity as a potential indicator of positive animal welfare. Animals. 2020 Jul;10(7):1211. doi.org/10.3390/ani10071211

60. Angiulli E, Pagliara V, Cioni C, Frabetti F, Pizzetti F, Alleva E, Toni M. Increase in environmental temperature affects exploratory behaviour, anxiety and social preference in Danio rerio. Sci Rep. 2020 Mar 25;10(1):1–2. doi.org/10.1038/s41598-020-62331-1.

61. Kim OH, Cho HJ, Han E, Hong TI, Ariyasiri K, Choi JH, Hwang KS, Jeong YM, Yang SY, Yu K, Park DS. Zebrafish knockout of Down syndrome gene, DYRK1A, shows social impairments relevant to autism. Mol Autism. 2017 Dec;8(1):1–4. doi.org/10.1186/s13229-017-0168-2

62. Kacprzak V, Patel NA, Riley E, Yu L, Yeh JR, Zhdanova IV. Dopaminergic control of anxiety in young and aged zebrafish. Pharmacol Biochem Behav. 2017 Jun 1;157:1–8.. doi.org/10.1016/j.pbb.2017.01.005

63. Maximino C, de Brito TM, da Silva Batista AW, Herculano AM, Morato S, Gouveia Jr A. Measuring anxiety in zebrafish: a critical review. Behav Brain Res. 2010 Dec 25;214(2):157–71. doi.org/10.1016/j.bbr.2010.05.031

64. Mathuru AS, Jesuthasan S. The medial habenula as a regulator of anxiety in adult zebrafish. Front Neural Circuits. 2013 May 27;7:99. doi.org/10.3389/fncir.2013.00099

65. Stewart A, Wu N, Cachat J, Hart P, Gaikwad S, Wong K, Utterback E, Gilder T, Kyzar E, Newman A, Carlos D, Chang K, Hook M, Rhymes C, Caffery M, Greenberg M, Zadina J, Kalueff AV. Pharmacological modulation of anxiety-like phenotypes in adult zebrafish behavioral models. Prog Neuropsychopharmacol Biol Psychiatry. 2011 Aug 1;35(6):1421–31. doi: 10.1016/j.pnpbp.2010.11.035. Epub 2010 Nov 29. PMID: 21122812.

66. Fontana BD, Cleal M, Parker MO. Female adult zebrafish (Danio rerio) show higher levels of anxiety-like behavior than males, but do not differ in learning and memory capacity. Eur J Neurosci. 2020 Jul;52(1):2604–13. doi.org/10.1111/ejn.14588

67. Roy T, Shukla R, Bhat A. Risk-Taking During Feeding: Between- and Within-Population Variation and Repeatability Across Contexts Among Wild Zebrafish. Zebrafish. 2017 Oct;14(5):393–403. doi: 10.1089/zeb.2017.1442.

68. Nadler LE, Killen SS, McCormick MI, Watson SA, Munday PL. Effect of elevated carbon dioxide on shoal familiarity and metabolism in a coral reef fish. Conserv Physiol. 2016 Jan 1;4(1):cow052. doi.org/10.1093/conphys/cow052

69. Mathuru AS, Kibat C, Cheong WF, Shui G, Wenk MR, Friedrich RW, Jesuthasan S. Chondroitin fragments are odorants that trigger fear behavior in fish. Curr Biol. 2012 Mar 20;22(6):538–44. doi.org/10.1016/j.cub.2012.01.061

70. Qin M, Wong A, Seguin D, Gerlai R. Induction of social behavior in zebrafish: live versus computer animated fish as stimuli. Zebrafish. 2014 Jun 1;11(3):185–97. doi.org/10.1089/zeb.2013.0969

71. Dreosti E, Lopes G, Kampff AR, Wilson SW. Development of social behavior in young zebrafish. Front Neural Circuits. 2015 Aug 18;9:39. doi.org/10.3389/fncir.2015.00039

72. Penalva A, Bedke J, Cook ES, Barrios JP, Bertram EP, Douglass AD. Establishment of the miniature fish species Danionella translucida as a genetically and optically tractable neuroscience model. bioRxiv. 2018 Jan 1:444026. doi.org/10.1101/444026

73. Vacher CM, O’Reilly JJ, Salzbank J, Lacaille H, Bakalar D, Sebaoui-Illoul S, Liere P, Clarkson-Paredes C, Sasaki T, Sathyanesan A, Kawasawa YI. Placental neurosteroids shape cerebellar development and social behaviour. bioRxiv. 2019 Jan 1:730150. doi.org/10.1101/730150

74. Buske C, Gerlai R. Shoaling develops with age in Zebrafish (Danio rerio). Prog Neuropsychopharmacol Biol Psychiatry. 2011 Aug 1;35(6):1409–15. doi.org/10.1016/j.pnpbp.2010.09.003

75. Fernandes Y, Gerlai R. Long-term behavioral changes in response to early developmental exposure to ethanol in zebrafish. Alcohol Clin Exp Res. 2009 Apr;33(4):601–9. doi.org/10.1111/j.1530-0277.2008.00874.x

76. Fernandes Y, Rampersad M, Gerlai R. Impairment of social behaviour persists two years after embryonic alcohol exposure in zebrafish: A model of fetal alcohol spectrum disorders. Behav Brain Res. 2015 Oct 1;292:102–8. doi.org/10.1016/j.bbr.2015.05.060

77. Kim OH, Cho HJ, Han E, Hong TI, Ariyasiri K, Choi JH, Hwang KS, Jeong YM, Yang SY, Yu K, Park DS. Zebrafish knockout of Down syndrome gene, DYRK1A, shows social impairments relevant to autism. Mol Autism. 2017 Dec;8(1):1–4. doi.org/10.1186/s13229-017-0168-2

78. Velazquez FN, Caputto BL, Boussin FD. c-Fos importance for brain development. Aging (Albany NY). 2015 Dec;7(12):1028. doi.org/10.18632/aging.100862

79. Yang P, Takahashi H, Murase M, Itoh M. Zebrafish behavior feature recognition using three-dimensional tracking and machine learning. Sci Rep. 2021 Jun 29;11(1):1–9. doi.org/10.1038/s41598-021-92854-0

80. Rosa LV, Costa FV, Canzian J, Borba JV, Quadros VA, Rosemberg DB. Three-and bi-dimensional analyses of the shoaling behavior in zebrafish: Influence of modulators of anxiety-like responses. Prog Neuropsychopharmacol Biol Psychiatry. 2020 Aug 30;102:109957. doi.org/10.1016/j.pnpbp.2020.109957

